# Translating genetics into tissue: inflammatory cytokine-producing TAMs and PD-L1 tumor expression as poor prognosis factors in cutaneous melanoma

**DOI:** 10.1101/2025.03.04.640087

**Authors:** Celia Barrio-Alonso, Alicia Nieto-Valle, Lucía Barandalla-Revilla, José Antonio Avilés-Izquierdo, Verónica Parra-Blanco, Paloma Sánchez-Mateos, Rafael Samaniego

## Abstract

Myeloid cells within tumor microenvironments exhibit significant heterogeneity and play a critical role in influencing clinical outcomes. In this study, we investigated the infiltration of various myeloid cell subtypes in a cohort of cutaneous melanomas, revealing no significant correlation between myeloid cell densities and the occurrence of distant metastasis. We further examined the phenotypic characteristics of primary melanoma tumor-associated macrophages (TAMs) utilizing the seven-phenotype classification recently proposed by Ma et al., derived from extensive pan-cancer single-cell RNA-sequencing studies. First, we analyzed the transcriptomic profile of TAMs isolated from stage IV metastasizing primary melanomas, alongside melanoma-conditioned monocytes cultured in vitro, both supporting the inflammatory cytokine-producing macrophage phenotype. Next, we employed multicolor fluorescence confocal microscopy, to assess the expression of TAM phenotype markers at the protein level in a cohort of primary melanoma samples. Notably, markers indicative of the inflammatory TAM phenotype, quantified at single-cell level, were significantly enriched in metastasizing tumors, demonstrating an independent correlation with shorter disease-free and overall survival (log-rank test, p< 0.0002). Additionally, our screening of phenotype markers expression revealed that PD-L1 positivity in tumor cells, rather than in TAMs, was associated with poor prognosis, highlighting a novel aspect of the immune landscape in cutaneous melanoma.

## Introduction

Myeloid cells play a pivotal role in the progression of melanoma, a highly aggressive form of skin cancer (Cheng et al., 2023, Trocchia et al., 2024). These cells are involved in both immune surveillance and modulation of the tumor microenvironment (TME), contributing either to tumor control or its advancement. The main subtypes that may infiltrate the TME are tumor-associated macrophages (TAMs), myeloid-derived suppressor cells (MDSCs), dendritic cells (DCs), granulocytes such as tumor-associated neutrophils (TANs) and eosinophils, and mastocytes (Mantovani et al., 2021). The prognostic significance of myeloid cell subtypes in melanoma underscores their complex roles, since their presence or abundance may significantly shape clinical outcomes and therapeutic responses. Although several studies have associated the abundance of certain myeloid cells with prognosis in human melanoma, many rely on blood markers or small cohorts of primary tumors (Ladanyi, 2015, Gartrell et al., 2018, Zilberg et al., 2025). Consequently, studies of primary tissues are yet to be conducted to determine their influence on prognosis and treatment response.

Macrophages progressively accumulate in most solid tumors, and an increased macrophage density is sometimes associated with poor prognosis in some types of cancer (Cassetta and Pollard, 2023). The different roles of TAMs come from their ability to adapt to the abundant environmental signals in the tumor microenvironment, showing protumoral or antitumoral functions (Glass and Natoli, 2016, Lopez-Janeiro et al., 2020). Although macrophage heterogeneity was initially described under two opposite phenotypes, M1 and M2, recent multi-omic analysis of macrophages in tumors like single-cell RNA-sequencing (scRNA-seq), metabolome and epigenome studies have shown that TAMs are highly heterogeneous, rendering potential distinct functional subsets (Ma et al., 2022, Mulder et al., 2021). Mulder et al. compiled 41 single-cell datasets from healthy and tumor tissues, defining numerous distinct clusters of monocytes and macrophages (MoMac-VERSE) (Mulder et al., 2021). Studies such as that of Ma et al., which propose seven TAM phenotypes, reflect a significant effort to consolidate and standardize the nomenclature of TAM diversity (Ma et al., 2022). Although single-cell transcriptomic analyses allow the identification and classification of distinct TAM clusters based on gene expression, the protein-level expression of many of these genes remains largely unknown, as not all messenger RNA is translated into protein. Therefore, bridging the gap between transcriptomic macrophage types and protein expression in tumor tissues is crucial for defining distinct TAM phenotypes or functional states that may be associated with prognosis or treatment response. Furthermore, characterization of these subsets at a protein-level, including their prognostic value in early-stage melanoma, may benefit immunotherapy strategies (Swetter et al., 2019).

Here we measured the density of myeloid cells in a cohort of paraffin-embedded primary melanomas and used protein expression data obtained through quantitative single-cell imaging to study the TAM heterogeneity and determine their prognostic potential in melanoma.

## Materials & methods

### RNA sequencing and transcriptomic analysis

Biopsied primary melanomas (n= 4) were homogenized and digested into single-cell suspensions (Tumor Dissociation Kit, Miltenyi Biotec), and TAMs were purified by magnetic cell sorting using CD14-microbeads (Miltenyi Biotec), as previously described (Samaniego et al., 2018, Gutierrez-Seijo et al., 2022). On the other hand, healthy donor monocytes were isolated and cocultured with BLM and A375 melanoma cell lines at a 1:2 ratio (melanoma:monocyte) for 24 hours and separated with CD14-microbeads. Once isolated (NucleoSpin RNA-purification kit, Macherey-Nagel Dueren, Germany), total RNA was processed and sequenced at BGI (https://www.bgitechsolutions.com) using the DNBseq-G400 platform. DEGs were assessed by using DESeq2 algorithm with parameters log_2_ (fold change) >2 and adjusted p_value <0.05. Both upregulated and downregulated DEGs were classified using the ‘enricher’ function of clusterProfiler R package (Wu et al., 2021) according to gene signatures published in three previous works (Ma et al., 2022, Mulder et al., 2021, Wei et al., 2024).

### Cohort study and selection criteria

Patient samples were collected following the approval of the Gregorio Marañón Hospital ethics committee, and written informed consent was obtained for each patient. A formalin-fixed and paraffin-embedded (FFPE) primary cutaneous melanoma cohort of 88 samples was used, all >2 mm Breslow thickness and a median follow-up of 81 months, excised between 1998 and 2015 in our institution. This cohort included 35 samples from patients who were disease-free for at least 10 years of follow-up (non-metastasizing primary melanomas) and 53 clinically aggressive samples developing distant metastasis (metastasizing primary melanomas, with 33/53 melanoma-related deaths). Pathological American Joint Committee on Cancer staging II–IV assessment was obtained through sentinel lymph node biopsy and distant metastasis evaluation by computed tomography at the time of diagnosis. Metastasizing and non-metastasizing primary tumors had comparable Breslow thickness (Mann–Whitney, p = 0.37). Six patients at stage IV were excluded from disease-free survival analysis, but not from overall survival. Due to the value of this cohort, the number of patients screened for each marker was maintained at the minimum statistically necessary, ranging from 15 in non-expressed markers to up to 74 in the case of inflammatory TAMs, which deserved the staining of four independent markers for their classification. Screened samples were randomly chosen from the 88 available patients.

### Multicolor fluorescence confocal microscopy

FFPE sections were deparaffinized and rehydrated, and antigens were retrieved by steaming in 10 mM sodium citrate buffer pH 9.0 (Dako, Glostrup, Denmark) for 7 minutes. Slides were blocked with 5 μg/ml human immunoglobulins solved in blocking serum-free medium (Dako) for 30 minutes and then incubated overnight at 4°C with 5– 10 μg/ml primary antibodies (supplementary Table S1), washed, and incubated with appropriate fluorescent secondary antibodies (Jackson Immunoresearch, West Grove, PA, USA) for 1 hour. Washes were performed in PBS containing 0.05% Tween-20. Single-cell quantification was performed for both density and protein expression at 3-5 20x fields. Mean fluorescence intensity (MFI) of proteins was obtained at manually depicted tumor cell nests or at automatically segmented CD68^+^, CD11b^+^ or CD66b^+^ cells using the ‘analyze particle’ plug-in of ImageJ2 software as previously shown (Barrio-Alonso et al., 2024, Gutierrez-Seijo et al., 2022). A glycerol-immersion ACS_APO_20x/NA 0.60 objective was used for quantification, and an ACS_APO_10x/NA 0.30 objective for the panoramic view (SPE confocal microscope, Leica). To allow suitable triple-staining combinations, we used a novel anti-Activin A antibody (R&D, AF338, antibody #2). VEGFA, CCL20 and TNF raw data from a previous survival study (Gutierrez-Seijo et al., 2021) are shown here for the first time to detect the putative inflammatory-TAM phenotype of the MoMac-VERSE. Non-expressed markers in melanoma TAMs were antibody validated in other human control tissues (Figure S1E). Ki67^+^ proliferating TAMs were so infrequent that could not be properly quantified by optical microscopy. Images are representative of markers co-staining, rather than density and/or clinical parameters.

### Statistical analyses

Kaplan–Meier curves were used to analyze the correlation with patient disease-free and overall survivals using Youden’s index to determine where the cutoff point was equally specific and sensitive. The Cox regression method (univariate and multivariate) was used to identify independent prognostic variables and Mann–Whitney tests to evaluate the association with clinicopathological features. Spearman correlation and log-rank analyses were also used in this study (GraphPad software, San Diego, CA, USA), as indicated; p < 0.05 was considered statistically significant.

### Data availability statements

Datasets can be found at https://www.ncbi.nlm.nih.gov/geo/query/acc.cgi?acc=GSE171277, hosted at Gene Expression Omnibus (GSE171277) (Gutiérrez Seijo A, 2021).

## Results

### Myeloid infiltration in primary melanomas

To investigate the infiltration of distinct myeloid cell populations in cutaneous melanoma, we quantified their density, both intratumorally and in the tumor periphery, in a cohort of paraffin-embedded primary melanomas (Figure 1A). Patients were classified as non-metastasizing or metastasizing, regarding subsequent development of metastasis during follow-up for 10 years. Quantification of markers for mastocytes (Triptase), TAMs (CD68), type 1 (CLEC9A) and type 2 (CD1c) DCs, TANs (CD66b), and eosinophils (Siglec-8) revealed no significant density differences associated with the metastatic behavior of the primary tumors (Figure 1B and C) or any other clinicopathological feature (supplementary Table S2). TAMs were by far the most abundant myeloid cells both inside and outside the tumors (macrophages represented the 90% of CD11b^+^ myeloid cells), while DCs and eosinophils were absent or scarce, located mainly in the periphery of the tumor. Density of TANs was, however, more erratic at both inside and outside regions of the tumors, being sometimes enriched in the periphery of non-metastasizing lesions (p= 0.02). Further characterization of TANs revealed that they do not express CD15, though more than half expressed CD33, a marker commonly associated with less differentiated MDSCs (Figure 1B and supplementary Figure S1A).

**Figure 1.**
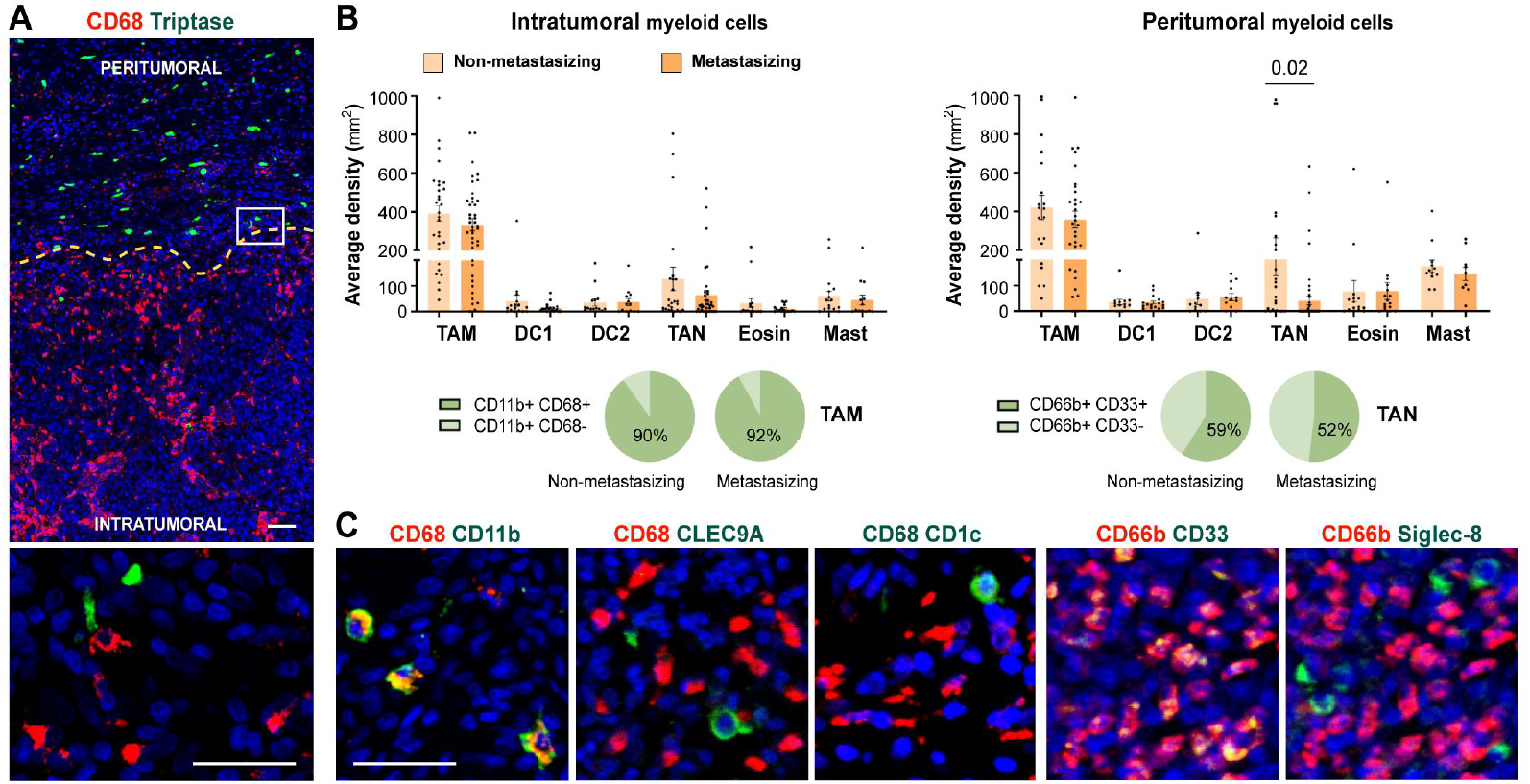
Myeloid cells density quantification in primary melanomas. **(A)** CD68^+^ TAMs (red) and Tryptase^+^ mast cells (green) enable primary melanoma delimitation of peritumoral and intratumoral areas. **(B)** Density quantification (mm^2^) of intratumoral and peritumoral myeloid cells comparing non-metastasizing and metastasizing primary melanomas. Percentages of CD11b^+^ CD68^+/-^ and CD66b^+^ CD33^+/-^ cells are shown, as indicated. **(C)** Representative images of CD68 (TAMs) and CD66b (TANs) in red, and CD11b (myeloid), CLEC9A (cDC1), CD1c (cDC2), or CD33 (MDSC) and Siglec-8 (eosinophil), in green. Dapi-stained nuclei, in blue. Scale bars, 50 μm.

### Primary melanoma macrophages and melanoma-conditioned monocytes acquire inflammatory cytokine-enriched TAM signature

Although the number of infiltrating TAMs does not vary between metastasizing and non-metastasizing tumors, crosstalk between TAMs and tumor cells (TCs) may induce the acquisition of gene expression profiles that may promote tumor progression (Gutierrez- Seijo et al., 2022). TAMs isolated from primary melanomas of stage IV patients (who had metastatic disease at the time of primary tumor removal), and therefore *bona fide* pro-metastatic TAMs, along with monocytes co-cultured for 24 hours with two different melanoma cell lines (BLM or A375), were subjected to RNA-sequencing to analyze their transcriptional profiles. Differentially expressed genes (DEGs) were subsequently analyzed for gene set enrichment using recently compiled TAM signatures by single-cell RNA-seq (scRNA-seq) (Ma et al., 2022, Mulder et al., 2021, Wei et al., 2024) (supplementary Figure S1B and Table S3). Both melanoma TAMs and melanoma-conditioned monocytes exhibited upregulated DEGs associated with cytokine-enriched inflammatory TAM phenotypes (named Inflam-TAM and Inflam-Cluster #15), alongside with a significant enrichment of the extracellular matrix-Cluster #13 of the MoMac-VERSE (Figure 2A and B). Conversely, both melanoma cell lines conditioned monocytes shared a significant downregulation of DEGs associated with the lipid-associated TAM phenotype (LA-TAM and LA-Cluster #3) (Figure 2A and B, and supplementary Figure S1C).

**Figure 2.**
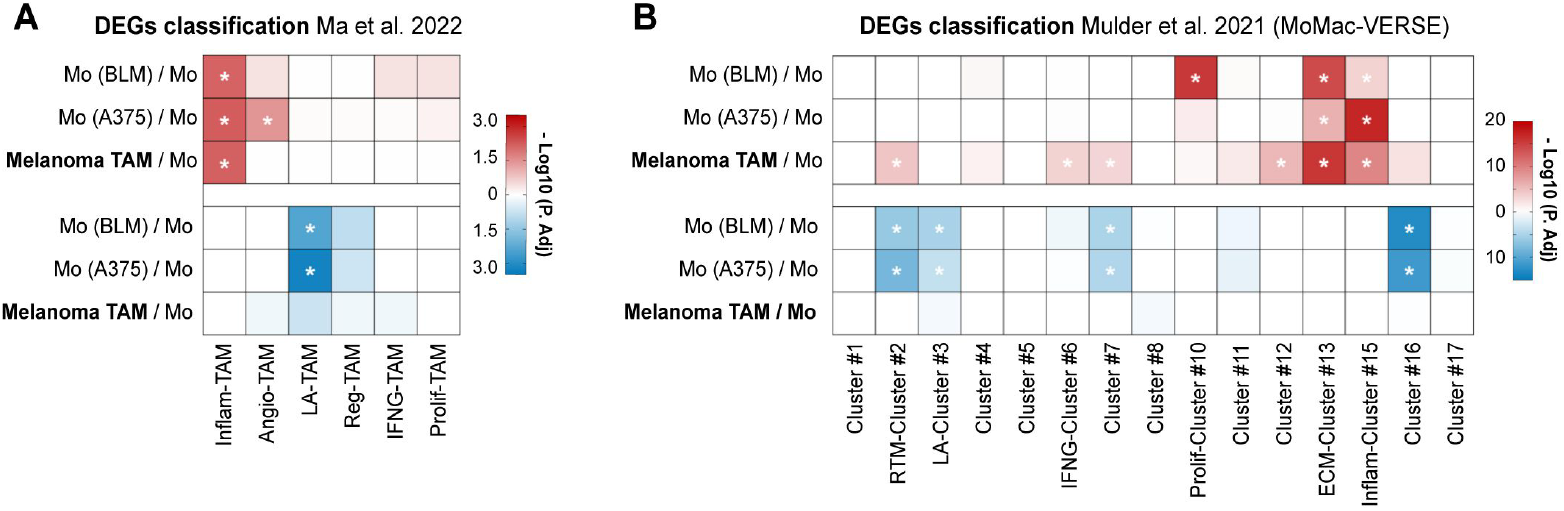
TAMs isolated from metastasizing primary melanomas show upregulated inflammatory gene signature. Differential expressed genes (DEGs) of TAMs isolated from stage IV primary melanomas (n= 4) and monocytes co-cultured with BLM and A375 melanoma cell lines (n= 3), compared to healthy donor monocytes, were classified according to gene sets proposed by Ma et al. **(A)** and Mulder et al. **(B)**. Heatmaps summarize the adjusted-p values for both upregulated (red) and downregulated (blue) DEG classification. *, P. adj <0.01.

### Single-cell protein expression of TAM markers proposed by scRNA-seq in primary melanomas and prognostic relevance of TC PD-L1

Since our transcriptomic analysis suggested that TAMs from stage IV metastasizing melanomas acquire a specific inflammatory profile among several other possible phenotypes, we designed a panel of antibodies to identify at the protein level the diverse macrophage types defined by Ma et al. at the transcriptional level. With selected antibodies, representative of each macrophage subset, we performed multiplex staining of our cohort of primary melanomas to explore their clinical significance (Figure 3A). The single-cell analysis of macrophage types was performed using multicolor fluorescence confocal microscopy, measuring the protein expression of the different markers in CD68^+^ TAMs, or in TCs if the studied marker was also expressed (Figure 3B-D). TAM expression of CCL8 and IDO was previously studied in the same cohort, showing no differences between non-metastasizing and metastasizing tumors (Gutierrez-Seijo et al., 2022, Barrio-Alonso et al., 2024), unlike the expression of VEGFA, CCL20, TNF and Activin A, which was significantly increased in metastasizing samples (p< 0.0001, Figure 3B). As VEGFA and CCL20 are shared markers by the inflammatory and the angiogenic macrophage subsets, we analyzed the expression of specific angiogenic markers HIF1A and FLT1. None of them showed detectable expression in melanoma TAMs, which was verified by proving the validity of the antibody in other human tissue macrophages (Figures 3B and supplementary Figure S1D and E). CXCL9, CXCL10, PD-L1, CD86, IL-10, TREM2, GPNMB, IL-4l1 and FOLR2 markers showed widespread expression by most melanoma TAMs, with no definition of a particular macrophage subset, and regardless of whether the tumor was metastasizing or not (Figure 3B and D and Supplementary Figure S1D). Interestingly, SPP1 was expressed by a small subset of macrophages, defining a unique type of TAM (supplementary Figure S1F).

**Figure 3.**
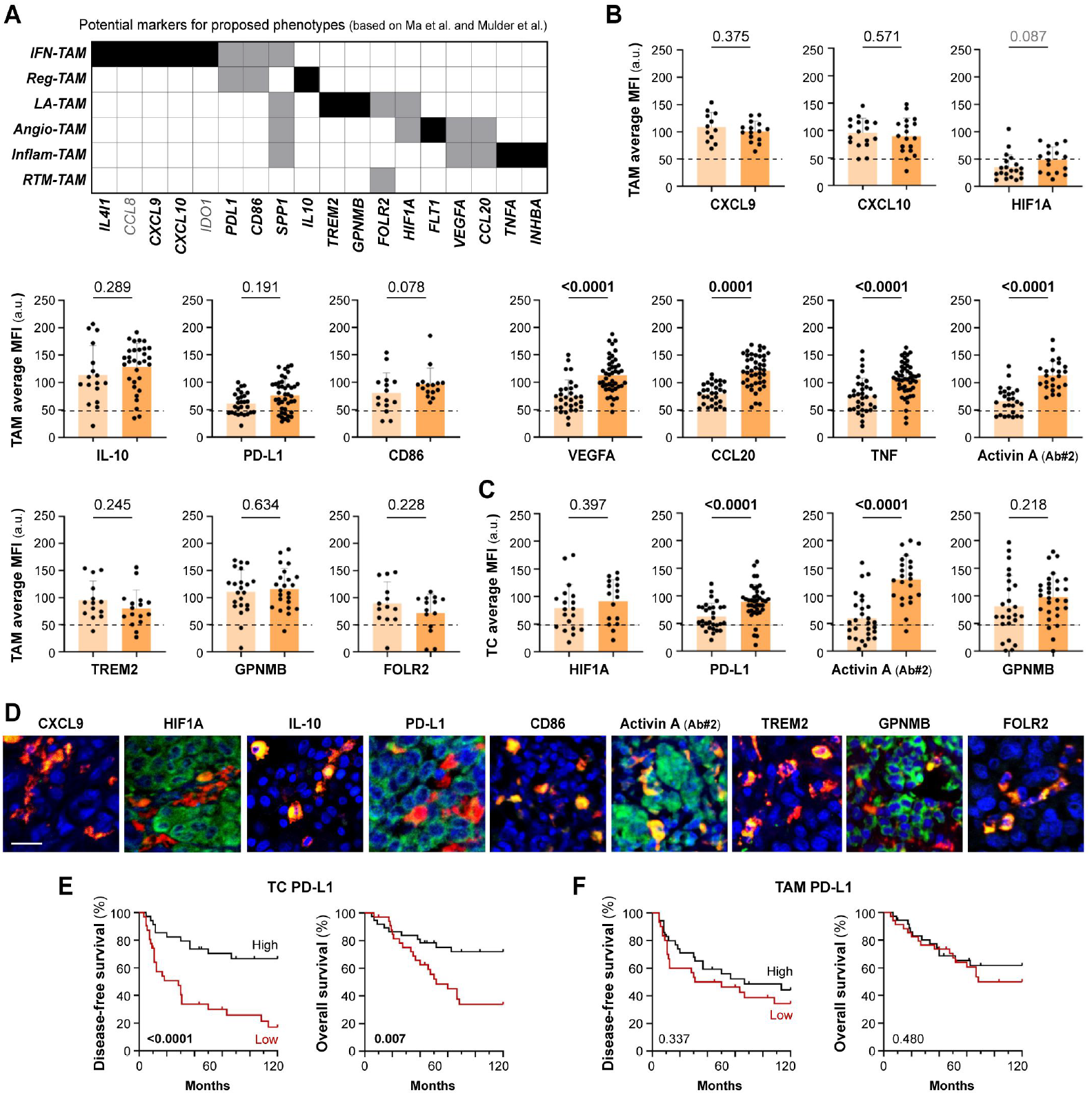
Protein expression of potential markers according to proposed TAM phenotypes. **(A)** Summary of potential markers for each proposed TAM phenotype (IFN-TAM, Reg-TAM, LA-TAM, Angio-TAM, Inflam-TAM and RTM-TAM), based on scRNA-seq pan-cancer compilations (Ma et al. and Mulder et al.) **(B, C)**. Single-cell protein quantification of CXCL9, CXCL10, HIF1A, IL-10, PD-L1, CD86, VEGFA, CCL20, TNF, Activin A, TREM2, GPNMB and FOLR2 at TAMs (B) or TCs (C), in non-metastasizing and metastasizing primary melanomas. Average mean fluorescence intensities (MFI) by patient are shown (arbitrary units, a.u.). Mann–Whitney p values, are indicated. **(D)**. Representative images of CD68^+^ TAMs (red) co-stained with indicated markers (green). Scale bar, 50 μm. **(E, F)**. Disease-free (E) and overall (F) survival Kaplan–Meier curves. Youden’s index was used to choose a cutoff point to classify primary melanomas as ‘low’ (red) or ‘high’ (black) for PD-L1 expression (n= 65) by TAMs (90 a.u.) and TCs (85 a.u.). Log-rank p values are shown.

Several of the markers selected to study TAM heterogeneity, including HIF1A, GMNMB, Activin A, and PD-L1, were also found to be expressed by cancer cells. Unlike PD-L1 expression by TAMs, PD-L1 expression by TC was found to be enriched in metastasizing melanomas (Figure 3C and D). We used the Kaplan-Meier method to assess the clinical relevance of PD-L1 expression by TCs in primary melanomas. Samples were stratified as ‘high’ or ‘low’ expression to calculate 10-year disease-free (DFS) and overall (OS) survival curves. Tumor cell PD-L1 expression correlated with shorter DFS (Log Rank, p< 0.0001) and OS (p= 0.007) in cutaneous melanoma (Figure 3E and F). Activin A expression by TCs and TAMs was previously reported to be associated with poor prognosis of melanoma patients (Gutierrez-Seijo et al., 2022), as confirmed here with a different anti-Activin A antibody (Figure 3D-F). Macrophage expression of Activin A at single cell level correlated positively with the expression of the inflammatory marker TNF (Spearman r= 0.52, p< 0.001), but not with PD-L1, FOLR2 or HIF1A which, as shown previously, were generally expressed by most TAMs (Supplementary Figure S1G). This suggests a notable commonality in TAM differentiation towards an inflammatory activation path in metastasizing samples, rather than enrichment in particular macrophage subtypes.

### Inflammatory cytokine-producing TAMs as activation biomarkers of poor prognosis in cutaneous melanoma

As both transcriptomic and single-cell protein analyses suggest the existence of an inflammatory cytokine-producing stage of pro-metastatic TAMs, we decided to perform a survival analysis using all four markers (Activin A, TNF, VEGFA, CCL20) simultaneously for sample stratification. Patients were stratified into two groups, those who highly expressed just 0-1 of the studied cytokines (non-inflammatory TAMs) *vs* 2-4 cytokine markers (inflammatory TAMs). Kaplan-Meier curves showed a strong correlation between inflammatory TAMs and DFS (p <0.0001) and OS (p= 0.0002) (Figure 4A). Furthermore, to determine whether the presence of inflammatory cytokine-producing TAMs and the expression of PD-L1 by TCs were independent prognostic factors, we performed a multivariate regression analysis including different clinicopathological features, such as age, gender, Breslow and staging. This analysis showed that the presence of inflammatory TAMs in tumors was an independent prognostic factor for DFS (p <0.0001) and OS (p= 0.006), as well as cancer cell PD-L1 (Table 1). Altogether, our findings suggest the potential of the inflammatory cytokine-producing path of TAM activation and the expression of Activin A or PD-L1 by TCs as clinically relevant biomarkers for patient stratification (Figure 4B and C).

**Table 1.**
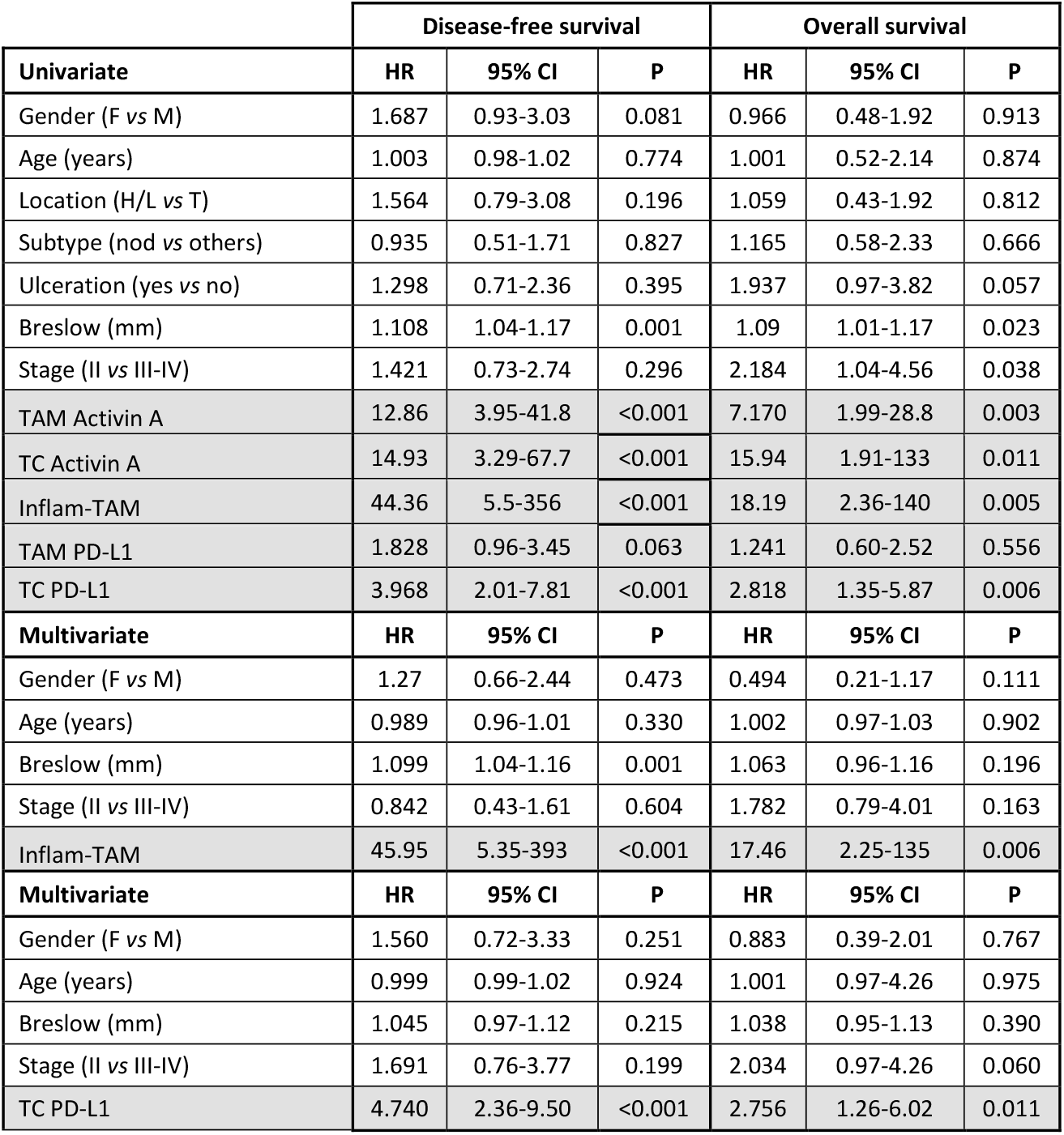
Univariate and multivariate Cox regression analyses for 10-year disease-free and overall survival. Abbreviations: CI, confidence interval; F, female; H/L, head/limb; HR, hazard ratio; M, male; Nod, nodular; T, trunk; TAM, tumor-associated macrophage; TC, tumor cell.

**Figure 4.**
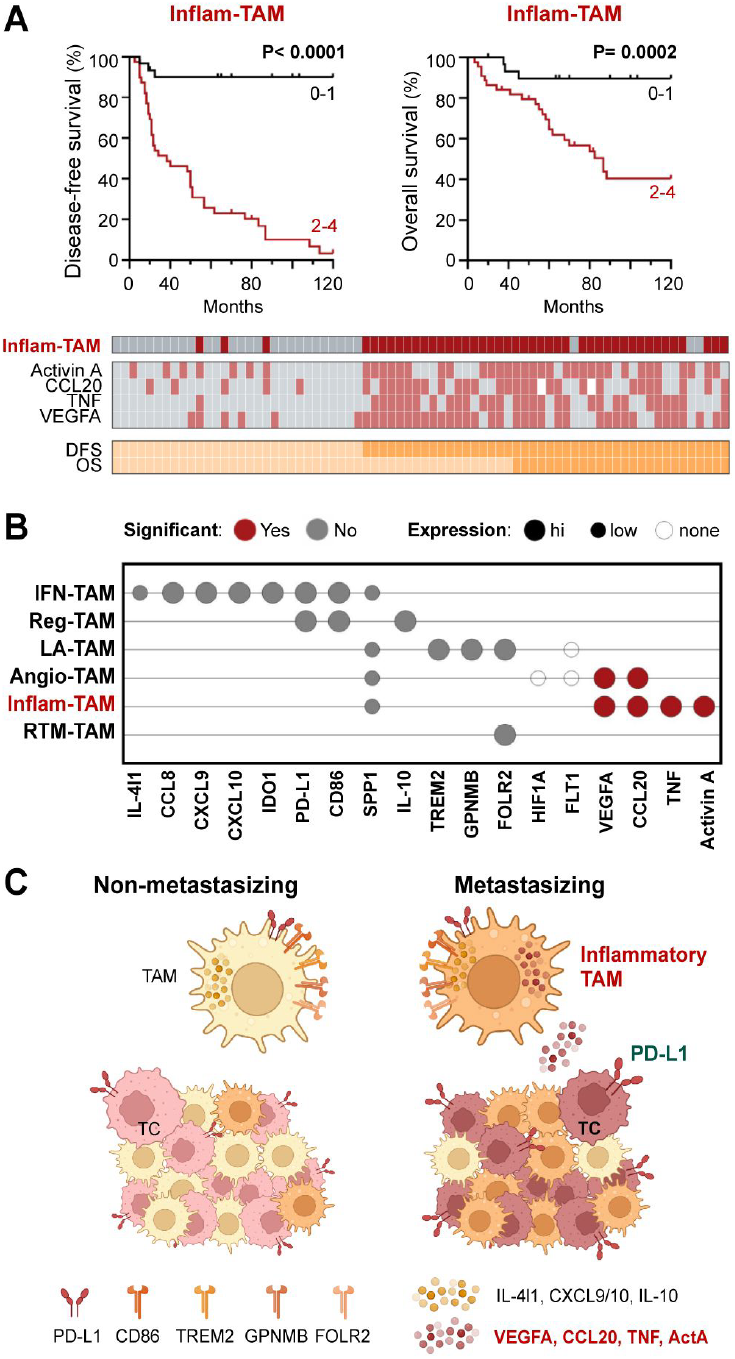
Inflammatory cytokine-producing TAMs (Inflam-TAMs) are associated with poor prognosis in cutaneous melanoma. **(A)** Disease-free and overall survival Kaplan– Meier curves. Primary melanomas were classified into two groups: expression of none-1 *vs* expression of 2-4 markers (Activin A, CCL20, TNF, VEGFA). Classification and expression of each marker in all analyzed primary tumors is shown. **(B)** Summary of the general (high, low, none) and relative (non-metastasizing vs metastasizing samples, Mann-Whitney p <0.01) expression of TAM markers at the protein level. **(C)** Resulting summarizing model, created with *BioRender*.*com*.

## Discussion

The innate immune system, as the first line of defense, is essential for controlling early tumor progression and initiating adaptive immune responses that ensure long-term tumor-specific immunity. Myeloid cells exhibit a remarkable plasticity which allows them to adopt either pro-tumoral or anti-tumoral roles depending on the surrounding cellular environment (Trocchia et al., 2024). Here the presence of myeloid populations was analyzed within and around tumors through a cohort of stage II-IV primary melanomas. TAMs constituted the main subset of myeloid cells in both compartments, while the rest of the subpopulations were a minority and preferentially located in the periphery of the tumor. TANs were an exception, since when present, they tended to form dense groups concentrated in specific regions of the tumor, mainly in the periphery of non-metastasizing lesions. This contrasts with a previous report in a collection of earlier-stage I-II melanomas, where the presence of intratumoral CD66b^+^ neutrophils correlated with worse prognosis (Jensen et al., 2012). Nevertheless, neutrophils have shown both anti- and pro-tumorigenic functions (Quail et al., 2022) and tumor staging might explain these differences.

The emergence of -omic technologies has significantly accelerated the detailed characterization of macrophages, proving that macrophage phenotypes are more complex than thought before, and that the M1-M2 classification is overly simplistic (Cheng et al., 2021, Mulder et al., 2021). Numerous studies have aimed to identify key molecular signatures to classify the heterogeneity of TAMs. Notably, two recent large-scale pan-cancer scRNA-seq analyses identified multiple potential subsets of TAMs. Although different nomenclatures were used in these two and other studies, similar molecular patterns were identified and a posterior review analysis unified the results and proposed that seven TAM subsets are conserved across nearly all cancer types (Ma et al., 2022). Despite not being single-cell, our bulk RNA-seq data obtained from TAMs isolated from already metastasized primary melanomas and from monocytes conditioned by melanoma cell lines, correlated significantly with the inflammatory cytokine-enriched TAM (inflam-TAM) phenotype (Mulder et al., 2021, Ma et al., 2022). Remarkably, several markers of this specific phenotype overlapped with our previously identified prognostic secretory signature (VEGFA, CCL20, TNF, Activin A), which proved to be triggered by melanoma cells via NFkB signaling pathway and sustained by the axis Activin A>Smad2/3 (Samaniego et al., 2018, Gutierrez-Seijo et al., 2022, Gutierrez-Seijo et al., 2021). Altogether our transcriptomic and single-TAM proteomic analyses suggest that the inflam-TAM phenotype from the MoMac-VERSE holds prognostic significance in melanoma and play most probably critical pro-metastatic and immune-suppressive roles.

Protein markers IL-4I1, CCL8, CXCL9, CXCL10 and IDO, that would allow the detection of interferon-primed TAMs (IFN-TAMs, a phenotype enriched in typical M1-like genes), were generally expressed in TAMs from both non-metastasizing and metastasizing tumors. Although none of these markers were associated with prognosis in our cohort of primary melanomas, the presence of this TAM phenotype has been previously associated with the response to immunotherapy in metastatic melanoma (Elewaut et al., 2025, Wei et al., 2024). Of note, in these studies interferon-primed TAMs are named inflammatory, which may lead to confusion with the inflammatory cytokine-enriched TAMs described by Ma et al. While multi-omics approaches have significantly advanced our understanding of cellular diversity, the absence of a unified nomenclature across studies, often assigning arbitrary names to cell clusters, risks creating misinterpretation and hindering cross-study comparisons.

Other markers expressed by both IFN-TAMs and immune regulatory TAMs (Reg-TAMs) are PD-L1 and CD86. Remarkably, despite no expression differences were found regarding TAM expression, PD-L1 expression by melanoma cells proved to be an independent prognostic biomarker, showing a strong negative correlation with both disease-free and overall survival. There are conflicting data from several studies regarding the prognostic value of PD-L1 expression in melanoma. A meta-analysis of these studies determined that its expression did not predict prognosis (Yang et al., 2020), however, the expression of this immune checkpoint protein was determined by classical non-quantitative IHC and distinction between TAMs and TCs was not considered. Furthermore, PD-L1 expression is also being studied as a predictive biomarker for immunotherapy in melanoma (Fiorentino et al., 2024), so quantitative image analysis together with separate analysis of TAMs and TCs should be considered from the point of view of treatment direction.

Lipid-associated TAM (LA-TAM) are characterized by the expression of lipid-related genes such as TREM2. This protein has been described as immunosuppressive in several types of cancer such as lung, colorectal and breast cancer (Katzenelenbogen et al., 2020, Molgora et al., 2020, Park et al., 2023). Another LA-TAM marker in blood-circulating monocytes, GPNMB, correlated with melanoma staging (Turrentine et al., 2014) and its expression by breast cancer TCs has been described as a prognostic indicator of recurrence (Rose et al., 2010). However, we did not detect expression differences for these proteins in our melanoma cohort, nor did we observe them for FOLR2, which was also widely expressed by the majority of macrophages, suggesting that this protein would not be an appropriate marker to identify the resident tumor-like TAM (RTM-TAM) subset in melanoma. In the same line, no differences were previously found for FOLR2 expression when comparing homeostatic and inflamed tissues, where the protein was broadly expressed by most of human tissue macrophages (Samaniego et al., 2014). Finally, despite pro-angiogenic TAMs (Angio-TAMs) have been well described in different tumors such as head and neck squamous cell carcinoma (Wu et al., 2024), their associated markers were barely detected in our melanoma cohort, suggesting that VEGFA expression might be associated with the inflammatory phenotype rather than the angiogenic subset.

Recent advances in single-cell RNA sequencing have provided a static view of TAM diversity, leading to new molecular definitions of macrophage states with potential prognostic and predictive significance in immuno-oncology. Although gene expression analysis yields valuable information, it does not consistently correlate with the presence of actual proteins within the tissue, so transcriptomic conclusions must be validated at the protein level (Ma et al., 2022, Jiang et al., 2020). Moreover, it remains critical to investigate the biological relevance and clinical impact of TAM subsets in larger patient cohorts, particularly in relation to their role in immunotherapy response.

Along with the heterogeneity of macrophage subsets, there are defined stages along four conserved paths of macrophage activation in tissues (Sanin et al., 2022). These activation stages include a “phagocytic” regulatory path, an “inflammatory” cytokine-producing path, an “oxidative stress” antimicrobial path, or a “remodeling” extracellular matrix (ECM) deposition path. Our results indicate that an inflammatory “cytokine-producing” stage of most TAMs, rather than enrichment in a particular macrophage subset is associated with poor patient prognosis in cutaneous melanoma.

## Acknowledgments

This work was supported by the Ministry of Science and Innovation PID2021-123507OB-I00 grant (PSM, RS) co-financed by ERDF/FEDER Funds from the European Commission, ‘A way of making Europe’; and by Cancer Research UK, FCAECC (GCB15152947MELE), and AIRC under the Accelerator Award program. CBA and ANV were financed by the Comunidad de Madrid YEI program.

## Author contributions

Conceptualization: CBA, ANV, PSM, RS; Data Curation: CBA, ANV, JAAI, VPB; Formal Analysis: CBA, ANV, LBR, RS; Funding acquisition: PSM, RS; Investigation: CBA, ANV, LBR, RS; Methodology: CBA, ANV, RS; Project administration: CBA, ANV, RS; Resources: JAAI, VPB, PSM, RS; Supervision: CBA, RS; Validation: CBA, ANV, RS; Writing-Original Draft Preparation: CBA, PSM, RS.

---

**Supplementary Table S1**. List of Antibodies used in this study.

**Supplementary Table S2**. Association of evaluated markers with clinicopathological features (Mann–Whitney).

**Supplementary Table S3**. Gene sets used for DEG classification.

**Supplementary Figure S1.**
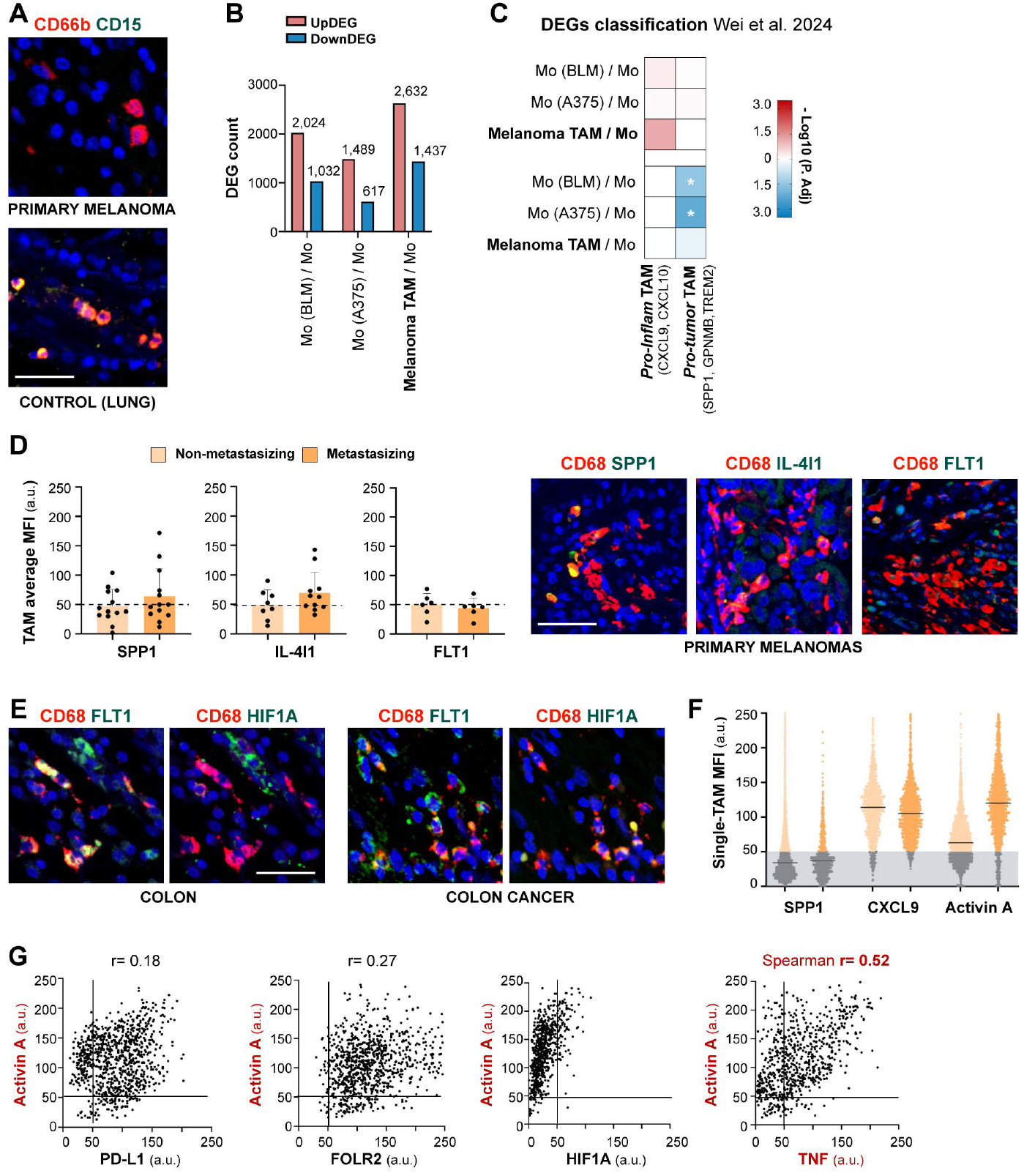
Melanoma associated macrophages phenotyping. **(A)** Representative FFPE human primary melanoma and healthy lung samples stained for neutrophil CD66b (red) and CD15 (green) markers. Scale bar, 50 μm. **(B)** Number of both upregulated (red) and downregulated (blue) De of macrophages isolated from primary melanoma tumors and monocytes co-cultured with melanoma cells lines. **(C)** Classification of DEGs of primary tumor TAMs and monocytes co-cultured with melanoma cell lines according to gene sets proposed by Wei et al. **(D)** TAM cell average MFIs for SPP1, IL-4I1 and FLT1 in primary melanomas (arbitrary units, a.u.) and representative primary melanomas stained for TAM CD68^+^ (red) and SPP1, IL-4I1 and FLT1 (green). **(E)** Representative FFPE healthy colon and colon cancer samples stained for CD68 (macrophages, red) and FLT1 and HIF1A (green). **(F)** Single-TAM quantification of SPP1, CXCL9 and Activin A (antibody #2) showing three distinct expression patterns: dichotomic, widespread, and differently expressed between metastasizing conditions, respectively. **(G)** Dot-plots showing single-cell expression of Activin A (antibody #2) vs PD-L1, FOLR2, HIF1A and TNF in TAMs (n= 3 metastasizing primary melanomas). Spearman r >0.4 is considered biologically relevant.

